# An Animal Model to Study *Klebsiella pneumoniae* Gastro-Intestinal Colonization and Host-to-Host Transmission

**DOI:** 10.1101/2020.02.04.934893

**Authors:** Taylor M. Young, Andrew S. Bray, Ravinder K. Nagpal, David L. Caudell, Hariom Yadav, M. Ammar Zafar

## Abstract

An important yet poorly understood facet in the life cycle of a successful pathogen is the host-to-host transmission. Hospital-acquired infections (HAI) resulting from the transmission of drug-resistant pathogens affect hundreds of millions of patients worldwide. *Klebsiella pneumoniae* (*Kpn*), a gram-negative bacterium, is notorious for causing HAI, with many of these infections difficult to treat as *Kpn* has become multi-drug resistant. Epidemiological studies suggest that *Kpn* host-to-host transmission requires close contact and generally occurs through the fecal-oral route. Herein, we describe a murine model that can be utilized to study mucosal (oropharynx and gastrointestinal [GI]) colonization, shedding within feces, and transmission of *Kpn* through the fecal-oral route. Using an oral route of inoculation, and fecal shedding as a marker for GI colonization, we show that *Kpn* can asymptomatically colonize the GI tract of immunocompetent mice, and modifies the host GI microbiota. Colonization density within the GI tract and levels of shedding in the feces differed among the clinical isolates tested. A hypervirulent *Kpn* isolate was able to translocate from the GI tract and cause hepatic infection that mimicked the route of human infection. Expression of the capsule was required for colonization and, in turn, robust shedding. Furthermore, *Kpn* carrier mice were able to transmit to uninfected cohabitating mice. Lastly, treatment with antibiotics led to changes in the host microbiota and development of a transient super-shedder phenotype, which enhanced transmission efficiency. Thus, this model can be used to determine the contribution of host and bacterial factors towards *Kpn* dissemination.

## Introduction

Host-to-host transmission of pathogens is the primary source of nosocomial infections, which are considered a serious threat to patient’s health and also a significant burden on the healthcare system (1, 2). Hospital-acquired infections (HAI) account for ∼100,000 deaths in the United States alone (3). A leading cause of these hospital-acquired infections and multiple outbreaks in hospitals around the world is *Klebsiella pneumoniae* (*K. pneumoniae*; *Kpn*), a member of the *Enterobacteriaceae* family that frequently causes pneumonia, bacteremia, pyogenic liver abscesses, and urinary tract infections (4), with most of these infections generally occuring in immunocompromised patients. With the rampant use of antibiotics *Kpn* isolates have become extensively drug-resistant, and some are now even considered pan-drug resistant, making the infections they cause extremely difficult to treat (5-7). For this reason, WHO lists *Klebsiella pneumoniae* as a critical pathogen for which new antibiotics and other therapies are urgently required to address this growing healthcare problem (8, 9). Further exacerbating treatment of *Kpn* infections is the recent identification of isolates termed “hypervirulent *K. pneumoniae*” (*hvKP*) that can cause disease, such as community-acquired pyogenic liver abscesses in healthy individuals (10-12). Patients recovering from *hvKP* infections often suffer from post-infectious sequelae that can lead to loss of limb or vision (13-15). These strains, originally isolated in the Pacific Rim, have since disseminated worldwide (10).

In the natural environment, the initial mucosal sites of colonization tend to be the oropharynx and the gastrointestinal (GI) tract (16, 17). These colonization events are generally asymptomatic (18). However, under certain circumstances, *Kpn* can gain access to other sterile sites in the host and cause disease. Epidemiological data suggest that many patients in hospitals are *Kpn* carriers in the GI tract, with a correlation between *Kpn* carriage and subsequent disease from the same isolate (19-21). Besides patients, hospital personnel can also be asymptomatic carriers of *Kpn*, and these silent carriers act as a reservoir from which *Kpn* can manifest disease within the same host or act as a source of transmission to a new host (18, 22-24).

Colonization resistance provided by the host microbiota plays a critical role in blocking colonization by pathogens. However, the use of antibiotics diminishes the microbial diversity in the GI tract, which potentially allows *Kpn* to readily colonize a host. Studies also show that antibiotic treatment of mice predisposes them to a “supershedder” state where they shed resident gut pathogens at a higher number, which enhances host-to-host transmission (25, 26). It is, however, unclear whether antibiotic treatment in a hospital setting contributes towards the increased transmission of drug-resistant *Kpn*.

Our understanding of the *Klebsiella pneumoniae*-associated disease-state comes mainly from animal models studying lung and urinary tract infection. While these studies have identified bacterial and host factors that contribute to *Kpn* virulence, there is very little mechanistic understanding of the gastrointestinal colonization and host-to-host transmission. Close contact, especially in a hospital setting, is thought to promote the spread of *Kpn* from an infected host to a naïve host. Transmission is thought to occur via the fecal-oral route, either through poor hygiene or contact with contaminated surfaces (fomites) (20, 22-24).

Here, we describe a novel murine model to allow for the study of *Kpn* GI colonization, shedding, and host-to-host transmission. Employing an oral route of *Kpn* inoculation in an inbred mouse population, we investigated *K. pneumoniae* gastric colonization and transmission. We demonstrate that *Kpn* can stably colonize the GI tract without treatment of antibiotics, and these mice stay persistently colonized and can transmit *Kpn* to cage-mates. Furthermore, antibiotic treatment of carrier mice induces gut dysbiosis and triggers a transient supershedder phenotype.

## Materials and Methods

### Ethics Statement

This study was conducted according to the guidelines outlined by National Science Foundation animal welfare requirements and the Public Health Service Policy on Humane Care and Use of Laboratory Animals (27). Wake Forest Baptist Medical Center IACUC oversees the welfare, well-being, and proper care and use of all vertebrate animals. The approved-protocol number for this project is A18-160.

### Bacterial growth conditions and strain construction

Strains used in the study are listed in Table 1. *K. pneumoniae* isolates were grown in Luria-Bertani (LB) Broth Lennox, with constant agitation at 37°C. For all mouse infections, an overnight culture of *Kpn* was spun down at ∼27000 *x g* for 15 minutes, and the resulting pellet resuspended in similar volume of 1X Phosphate-Buffered Saline (PBS). To obtain desired density for mouse infections (10^6^ CFU/100µl), the bacterial suspension in PBS was diluted into a 2% sucrose-PBS solution. Ten-fold serial dilutions were plated on selective media (LB-Agar with antibiotic) and incubated at 30°C overnight for quantitative culture. LB plates containing antibiotics were streptomycin (str, 500 µg ml), chloramphenicol (50 µg/ml), ampicillin (25 µg/ml), apramycin (50 µg/ml), spectinomycin (30 µg/ml) and rifampin (30 µg/ml).

The *wzi* gene codes for a conserved outer membrane protein involved in the attachment of capsular polysaccharide to the outer membrane. Sequence polymorphism in the *wzi* gene has been used to identify and characterize different isolates. *Kpn* AZ10 (*wzi* 372) an antibiotic sensitive isolate was made str-resistant as described, subsequently mouse GI passaged and named AZ99 (28, 29). To construct the *fimH* mutant in the appropriate genetic background, PCR was carried out using Q5 polymerase (NEB) with AZ101 genomic DNA as template, and primers *fimH upstream* (GGCGGTGATTAACGTCACCT) and *fimH downstream* (GATAGAGCAGCGTTTGCCAC), which give at least 500bp homology on either end of the transposon cassette. The PCR product was purified using the Qiagen MinElute kit. Lambda Red mutagenesis was carried out as described previously (30), and cells were recovered in Super Optimal Broth with Catabolite repression (SOC) media at 30°C with shaking overnight. Recovered bacteria were plated on selective LB agar containing chloramphenicol (50 µg/ml). Single colonies were purified and the mutation was confirmed by PCR.

To determine *fim* promoter orientation, PCR was carried out using either *in vitro* samples from LB broth, single colonies from LB plates, or *in vivo* samples (fecal pellets) from mice infected with *Kpn*. Broth culture was spun down as described above and resuspended in equal volume dH_2_O and boiled for 5 minutes. Also, a single colony was resuspended in 10 µl of dH_2_O and boiled for 5 minutes. DNA was isolated from fecal pellets (100 mg) using Quick-DNA™ Fecal/Soil Microbe Microprep (Zymo Research). 5 µl of sample was used in PCR with OneTaq polymerase (NEB) with primers Cas168 GGGACAGATACGCGTTTGAT and Cas169 GCCTAACTGAACGGTTTGA as described previously (31). Purified PCR product was digested with restriction enzyme *HinFI* (NEB) for 1 hour and resolved on 1.2% agarose gel. As established previously, the “off orientation” of the *fim* promoter results in product bands of 496bp and 321bp, whereas the “on orientation” results in bands of 605bp and 212bp (31).

### Mouse Infections for Colonization and Shedding

Colonies of C57BL/6J (SPF) mice obtained from Jackson Laboratory (Bar Harbor, ME) were bred and maintained in a standard animal facility at Biotech Place, Wake Forest Baptist Medical Center. All animal work was done according to the guidelines provided by the American Association for Laboratory Animal Science (AALAS) {Worlein, 2011 #87} and with the approval of the Wake Forest Baptist Medical Center Institutional Animal Care and Use Committee (IACUC). 5-7 week-old mice were infected and monitored through the course of the experiments. Food was removed from mice ∼6 hours prior to inoculation. Mice were fed ∼10^6^ CFU/100 µl of *K. pneumonia* in two 50 µl 2% sucrose-PBS doses an hour apart, from a pipette tip. Immediately afterwards, food was returned to mice.

To quantify daily bacterial shedding, mice were removed from their housing and placed into isolation containers. Fecal pellets (∼0.02 g, approximately 2 pellets) were collected and placed into a 2 ml screwcap tube (Fisherbrand, 02-682-558) along with at least 2 glass beads (BioSpec, 11079127). Samples were diluted 1:10 in PBS (weight:volume). A Bead mill 24 (Fisherbrand) was used to homogenize the fecal pellets (2.1 power setting, 1 min). Afterwards, the tubes were spun in a mini centrifuge (Thermo Scientific, MySpin 6) to pellet out larger debris. Ten-fold serial dilutions were plated from the supernatant on appropriate antibiotic plates and incubated overnight at 30°C. Bacterial shedding was calculated in CFUs per gram of feces. The limit of detection was 100 CFU/g. Each mouse was uniquely marked so that the fecal shedding of each individual mouse could be tracked for the duration of the experiment.

To trigger an antibiotic-dependent *Kpn* supershedder phenotype, mice were infected orally with a str-resistant *Kpn* as described above. Four to five days post-inoculation (p.i) mice were gavaged with streptomycin (5 mg/200 µl) either once or on three consecutive days, and daily shedding monitored post-antibiotic treatment. To determine effect of neomycin treatment before *Kpn* infection, mice were gavaged with a single dose (5 mg/200 µl) 24 hours before inoculation with the MKP103 a derivative of KPNIH1 isolate with a deletion of the KPC-3 carbapenemase-encoding gene (32). Bacterial counts were enumerated from fecal pellets as described above. To determine the role of continuous antibiotic treatment on supershedder phenotype, the drinking water was replaced with water containing 1g/L ampicillin 24 hours before infection; mice were maintained on ampicillin-water for 10 day p.i. after which they were placed on regular water until the end of the experiment. *Kpn* fecal shedding was assessed up to 20 days post-infection, and quantified as described above.

For competition experiments, mice were infected with a 1:1 mixture of AZ94 and the mutant of interest. Fecal shedding of both strains was assessed as described above. Fecal homogenates were plated on both apramycin (50 µg/ml) and str (500 ug/ml) LB agar. The competitive index (CI) was calculated as described previously (33), using the following equation:

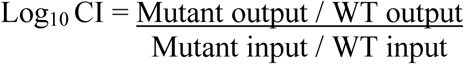

A value of 0 would suggest that neither strain has an advantage. A value >1 would suggest that the mutant has competitive advantage, whereas a value <1 would indicate the WT has the advantage.

To determine colonization density in the GI tract, ileum, cecum, and colon were removed under sterile conditions immediately following CO_2_ (2 liters/min, 5 min) euthanasia of animals and subsequent cardiac puncture. The cecum, proximal colon, and a span of the terminal section of the ileum equal in length to the colon were removed from each animal. Organs were weighed and placed into individual 2 ml screwcap tubes (Fisherbrand, 02-682-558) with at least 2 glass beads (BioSpec Products, 11079127). Samples were diluted 1:10 in PBS (weight:volume) and were homogenized and plated as described above. The limit of detection was 100 CFU/g.

To determine colonization density in the kidney, liver and spleen, organs were removed under sterile conditions immediately following euthanasia as described above. Organs were weighed and placed 15 ml conical tubes. For kidney and liver equal weight to volume PBS was added, and samples were homogenized using PowerGen 700 (Power setting 2 for 30 seconds), whereas for spleen, ten times the volume of 1X PBS was added to the weight of the organ and homogenized as above. The samples were plated as described. The limit of detection of kidney and liver was 33 CFU/ml and for spleen 100 CFU/ml.

Oropharyngeal lavage was carried out with 200 µl of sterile PBS from a gavage needle inserted into the esophagus. The esophagus was exposed and cut transversely. A gavage needle, attached to a prefilled insulin syringe (BD) with 1X PBS was then inserted into the cut esophagus, and PBS collected from the mouth. The collected lavage was serially diluted and plated on appropriate antibiotic plates and incubated overnight at 30°C. The limit of detection for oral lavage was 33 CFU/ml.

### Transmission Studies

For 4:1 and 1:4 transmission experiments C57BL/6J index mice (n=1 or n=4) at 5-7 weeks of age are infected with *Kpn* as described above, and shedding is collected daily to determine colonization density of the GI tract. On day 4 p.i., contact mice (n=4 or n=1) were introduced to cages with the index mice. Fecal shedding of index and contact mice was collected and quantified for at least 6 days post-cohousing, for a total of 10 days for index mice and 6 days for contact mice. On day 10, the mice were euthanized as described above and the ileum, cecum, colon, and oral lavages of all mice were processed as described above to determine colonization density of *Kpn*.

In 1:4 transmission experiments, in which the administration of a single dose of antibiotic was assessed, index mice were infected with a str-resistant *Kpn* and fecal shedding of *Kpn* was quantified for 4 days p.i. On day 5 p.i, index mice were treated with streptomycin (5 mg/200 µl) via gavage and then co-housed with contact mice. Fecal shedding was collected daily from index and contact mice. In a continuous antibiotic challenge transmission study, an index mouse was put on water containing ampicillin (1 g/L) 24 hours before infection. The contact mice were placed on water containing ampicillin (1 g/L) 24 hours before introduction of the index mouse. Once co-housed, daily shedding was collected from both index and contact mice to determine if any transmission events occurred.

To confirm that host-to-host transmission events occur through the fecal oral route, a metabolic cage (Tecniplast Cat. # 3700M022) was used. *Kpn* infections were carried out as described above. 4 days post-infection, a contact mouse was introduced into the metabolic cage and fecal shedding collected from both index and contact mice to determine transmission frequency.

### Histology

Mice were infected with either PBS (vehicle-only control) or a KPPR1S isolate. A subset of *Kpn*-infected mice were gavaged with streptomycin (single treatment; 5 mg/200 µl) to induce the super-shedder state at 5 days p.i. As a positive control, mice were put on 3% w/v Dextran Sodium Sulfate (DSS) molecular weight 50,000 in their drinking water *ad libitum* for 7 days. All the mice were euthanized at day 7 post initial treatment or infection. 2.5 cm of colon immediately distal to the cecum was collected, washed with 1x PBS, and prepared using the Swiss roll method. Afterwards the sample was preserved in 1:10 Formalin (Fisherbrand, 305-510), and after 24 hours transferred to 70% ethanol. The samples were embedded in paraffin before being sectioned, mounted, and stained with hematoxylin and eosin (H & E). The resulting slides were scored by the Wake Forest Baptist Medical Center Pathology department.

Liver was collected under sterile conditions from either mock infected mice or from orally infected mice with hvKP1, and in extremis. Liver samples were cut in to sections about 6.5mm, and placed in 10% formalin (10 parts formalin to 1 part tissue). After 24-48 hours the samples were transferred to 70% Ethanol and stored at 4°C till they were further processed for H & E and Gram staining, with scoring carried out as described above.

### Fecal microbiome Analysis

Fecal microbiome was examined according to previously described methods (34-36). Briefly, genomic DNA from 200 mg feces was extracted using MoBio Powerfecal DNA kit (Qiagen, Valencia, CA) per manufacturer’s instructions. Amplicon PCR of the V4 hypervariable region of the 16S rDNA gene was performed using the universal primers 515F (barcoded) and 806R according to the Earth Microbiome Project protocol (PMID: 22402401). The amplicons were purified using AMPure^®^ magnetic beads (Agencourt), and the products quantified with Qubit-3 fluorimeter (InVitrogen). The final amplicon library was generated as previously described (37). Equimolar pooled library was sequenced on an Illumina MiSeq platform using 2×300bp reagent kit (Miseq reagent kit v3; Illumina Inc.) for paired-end sequencing. The sequencing quality control was done with on-board Miseq Control Software and Miseq Reporter (Illumina Inc.) and the obtained sequences were de-multiplexed, quality-filtered, clustered and analyzed using QIIME software package (34, 35, 38, 39). Taxonomy classification was performed within QIIME based on 97% sequence similarity to the Greengenes database (38). Alpha-diversity and bacterial proportions were compared using Kruskal-Wallis test followed by pair-wise Mann-Whitney test. Linear discriminatory analysis (LDA) effect size (LEfSe) was applied to identify discriminative features (unique bacterial taxa) that drive differences at different time-points or in different groups (40). Hierarchical clustering and heat-maps depicting the patterns of abundance were constructed within ‘R’ statistical software package (version 3.6.0; https://www.r-project.org/) using the ‘heatmap.2’ and “ggplots” packages.

### Statistical analysis

All statistical analyses were performed using GraphPad Prism 8.0 (GraphPad Software, Inc., San Diego, CA). Unless otherwise specified, differences were determined using the Mann-Whitney *U* test (comparing two groups) or the Kruskal-Wallis test with Dunn’s post-analysis (comparing multiple groups).

## Results

### Establishing *Klebsiella pneumoniae* colonization in the murine intestinal tract

We sought to establish a GI model of *Klebsiella pneumoniae* colonization that would mimic natural colonization in a host. Because of the difficulty in establishing *Kpn* GI colonization through gavage treatment, previous studies used antibiotic pre-treatment to disrupt the host microbiota and allow for *Kpn* colonization via the gavage method (41-43). We first tested the ability of *Kpn* to colonize the GI tract by giving adult mice doses ranging from 10^5^-10^9^ CFU/100µl, without antibiotic treatment so as not to disrupt the host microbiota (**Fig. 1A**). However instead of a gavage treatment, mice were infected orally by pipette feeding to simulate the natural route of infection (44). We used a *Kpn* clinical isolate KPPR1S that has been used extensively to model *Kpn*-associated disease-state in mice. The streptomycin- and rifampin-resistance of KPPR1S allowed us to enumerate the bacteria in the fecal pellets on selective plates. As observed in **Fig. 1B-C**, *Kpn* colonized the GI tract and was shed robustly in the feces of mice with doses above 10^5^ CFU. A dose < 10^5^ CFU did not result in the establishment of colonization (Threshold for detection 100 CFU), suggesting a minimum dose of 10^5^ is required (Data not shown). Based upon these results, we chose 10^6^ CFU, as the minimum dose required to establish *Kpn* colonization. Based on our preliminary studies that suggest poor *Kpn* fecal shedding correlates with reduced GI colonization, we used daily fecal shedding as a substitute for colonization density in the GI tract. Next, we determined how long *Kpn* colonizes the mouse GI tract. We followed *Kpn* shedding in feces of infected mice for either 15 or 30 days p.i and observed that *Kpn* was shed at similar levels throughout the study **(Fig. 1D-E)**. Furthermore, our results showed that *Kpn* colonizes the mucosal surface of the oropharynx (**Fig. 1F**). Taken together, our data suggest that, when introduced by the oral route, *Kpn* colonizes the mucosal surface of the oropharynx, can establish and persist in the GI tract, and is shed robustly in the feces.

**Figure 1.**
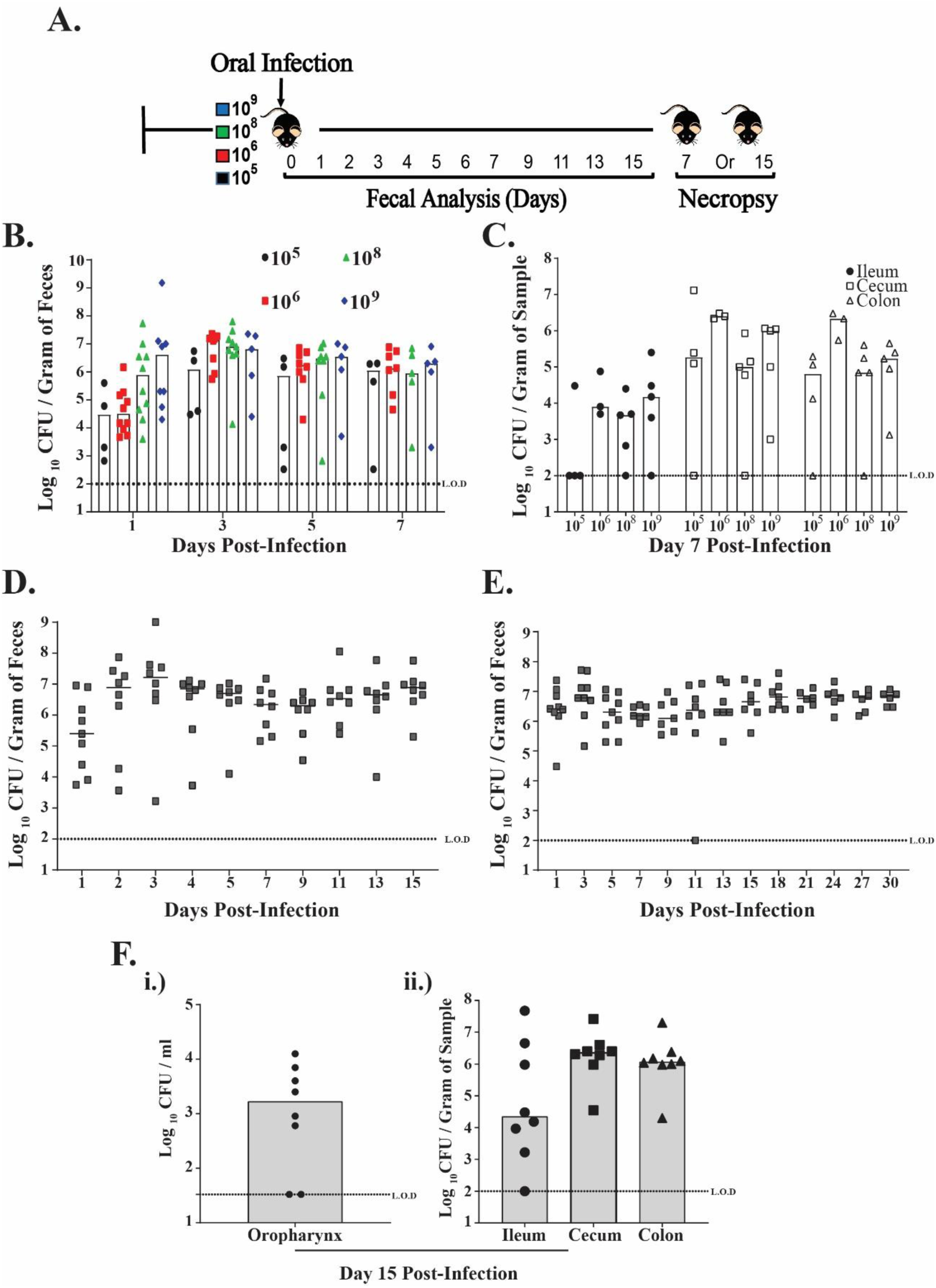
*K. pneumoniae* colonizes the gastro-intestinal tract of mice. **(A)** A schematic representation of C57BL/6J mice orally infected with *K. pneumoniae* KPPR1S isolate (10^5^, 10^6^, 10^8^ and 10^9^ CFU). **(B-C)** shows quantified daily shedding results and colonization density in the intestinal tract (ileum, cecum and colon) at day 7 post-inoculation from mice given different *Kpn* doses. **(D-E)** Fecal shedding data collected from mice given 10^6^ CFU of KPPR1S and followed for up to either 15 days or 30 days post-infection. **(F) (i, ii)** Colonization density of KPPR1S isolate in the oropharynx and lower GI tract of mice 15 days post-inoculation. Bars indicate the median values. L.O.D, limit of detection.

A hallmark of *K. pneumoniae* isolates is their genetic heterogeneity, which affects their ability in causing disease (45). Thus, we determined whether *Kpn* genetic plasticity also contributes to GI colonization. We tested the ability of a set of genetically diverse clinical *Kpn* isolates to colonize the GI tract of mice. For analysis we chose MKP103 a derivative of KPNIH1, which was the cause of an outbreak at NIH Clinical Center, hvKP1, a hypervirulent human isolate, and AZ99, a human fecal isolate. All three strains showed varying colonization density of the murine GI tract, with the hvKP1 shedding at a similar level to KPPR1S (**Fig. 2A**). Surprisingly, the MKP103 isolate colonized poorly, with mice generally clearing it from their GI tract by day 5 p.i. As observed through fecal shedding, the human fecal isolate AZ99 consistently colonized the GI tract albeit at a lower density in comparison to KPPR1S. Moreover, mice colonized with hvKP1 had a high mortality rate (**Fig. 2C**). Hypervirulent isolates are notorious for causing pyogenic liver abscesses (PLA) (11). As shown in **Fig. 2D** mice that succumbed after oral inoculation with the hvKP1 isolate, were colonized at a high density in the liver, kidney and spleen with the same isolate. Moreover, these mice appeared to have developed liver abscesses **(Fig. 3A)**, which H&E and Gram staining confirmed to contain necrotic tissue, inflammatory cells, and gram negative bacteria **(Fig. 3B-D)**. Thus, our model mimics human disease dynamics, where a hypervirulent isolate (hvKP1) is able to translocate from the GI tract to other sterile sites, and cause the development of the disease state.

**Figure 2.**
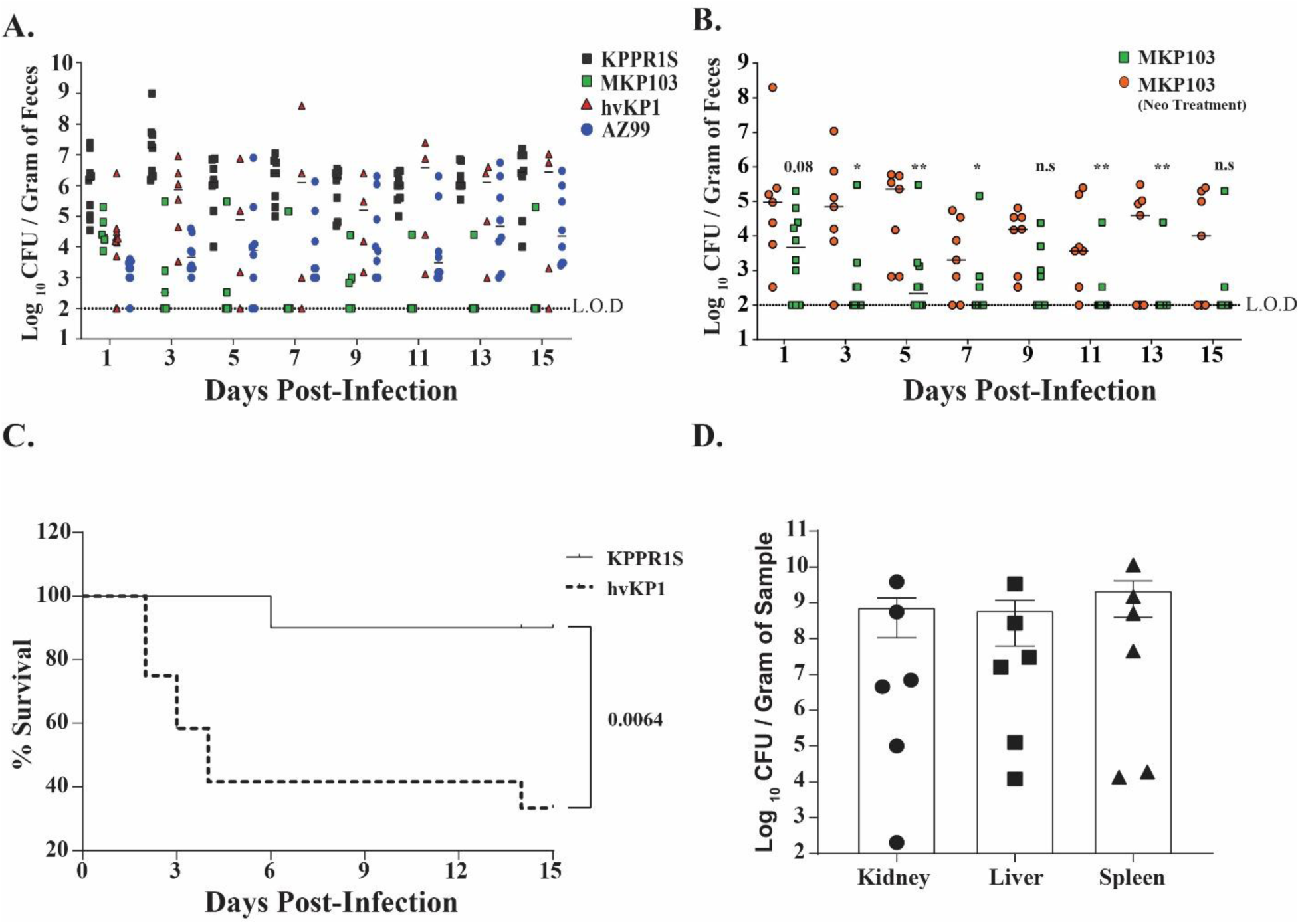
Differences among *K. pneumoniae* clinical isolates in fecal shedding levels and virulence. **(A.)** Mice were infected with the indicated *K. pneumoniae* isolate with daily shedding values shown. **(B.)** Comparison between mice infected with *K. pneumoniae* strain MKP103 with or without pre-treatment of antibiotic neomycin (5 mg/ 200 µl). Symbols represent shedding values obtained from a single mouse on a given day. The bar represents the median value. Dashed line represents the limit of detect (L.O.D). **(C.)** Survival of mice infected gastro-intestinally with *K. pneumoniae* isolate KPPR1S (n=10) or hvKP1 (n=10) over 15 days. An in extremis state or death was scored as non-survival. Log-rank (Mantel-Cox) test performed to determine statistical differences. **, *P <*0.01 for KPPR1S compared to hvKP1. **(D.)** Bar graph showing mean colonization density of hypervirulent isolate hvKP1 in the kidney, liver and spleen from mice that were initially infected orally with S.E.M bars. Each data point represents colonization density in a specific organ from a specific mouse. Organs were harvested from mice that displayed an in extremis state. Limit of detection of liver and kidney was 33 CFU/ml and 100 CFU/ml for spleen.

**Figure 3.**
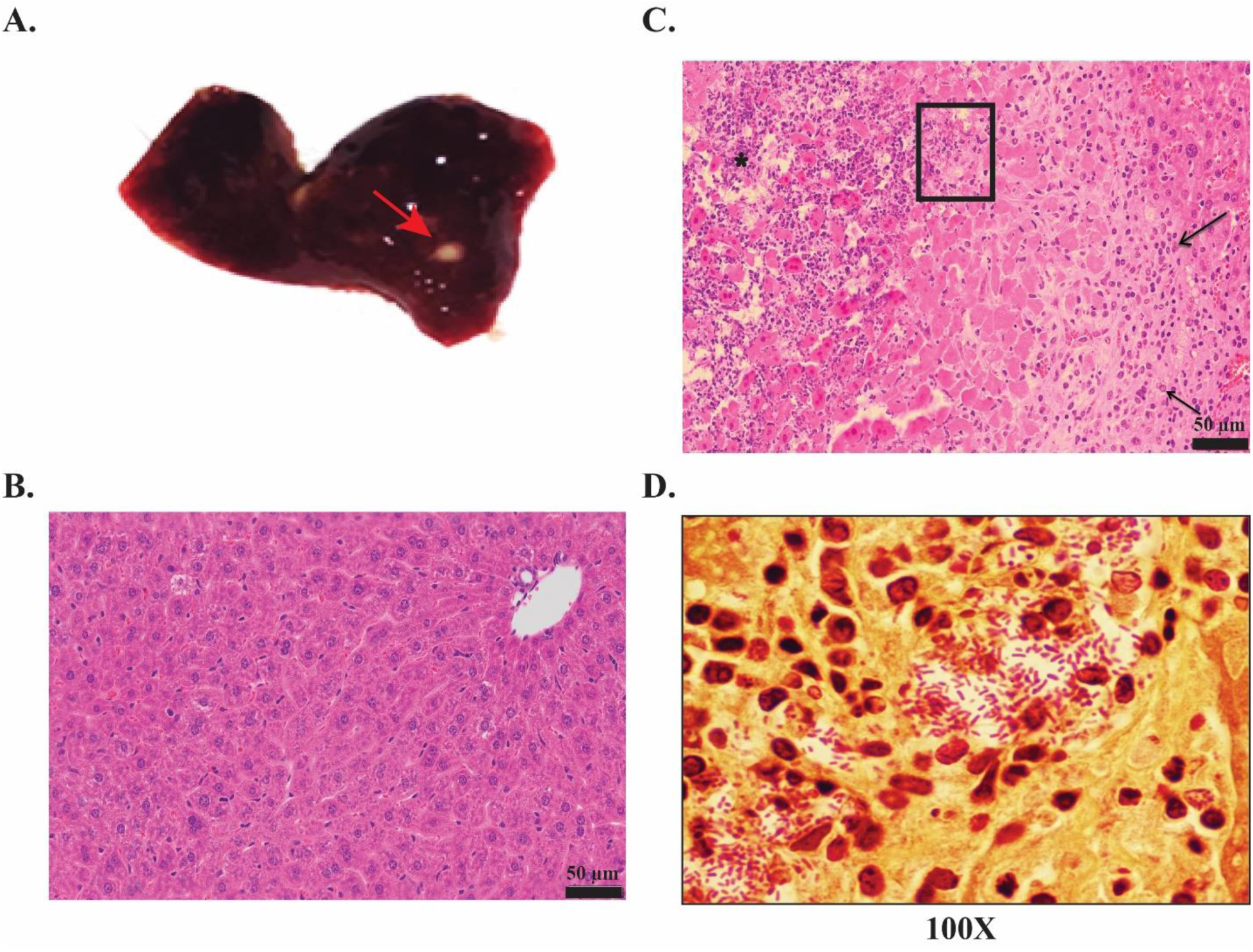
Representative *K. pneumoniae* liver abscesses (Red Arrows) from mice that succumbed to infection from hvKP1 isolate that originally colonized their GI tract **(A.)**. Liver tissue samples stained with hematoxylin and eosin staining. Shown here are liver tissue sample at 20X resolution from either uninfected mice **(B.)** showing normal heptaocytes or from infected mice **(C.)** that contains a large region of coagulative and lytic necrosis with hepatocellular disassociation including central infiltrates of neutrophils (*) with few scattered mononuclear inflammatory cells. The peripheral aspect of the lesion contains numerous lymphocytes, macrophages and new scattered neutrophils (arrows). (D) 100X oil immersion image of the boxed area from C showing Gram negative bacteria present within the necrotic liver tissue.

### Antibiotic treatment leads to the development of the *Kpn* supershedder phenotype

Given that the fecal-oral route of transmission in a hospital setting is considered a significant cause of nosocomial infections (46, 47) it was surprising that the MKP103 isolate failed to colonize the GI tract of mice (**Fig. 2A**). However, as many of the patients that acquired MKP103 in the GI tract were on antibiotics (20), we considered whether the use of antibiotics would affect the ability of this isolate to colonize the GI tract. Moreover, high use of antibiotics in a health-care setting correlates with *Klebsiella pneumoniae* infections (20). Therefore mice were gavaged with neomycin to reduce the colonization resistance by the host GI microbiota, and then infected with MKP103 to determine whether antibiotic treatment positively affected its ability to colonize. As shown in **Fig. 2B** antibiotic pre-treatment of mice allowed MKP103 to colonize and persist within the infected host GI tract up to 15 days post-infection.

Our results show that antibiotic treatment allows *Kpn* isolate (MKP103) that colonizes poorly to establish itself in the GI tract. However, whether antibiotic treatment affects colonization density of isolates that colonize robustly without requiring antibiotic treatment remains unknown. We determined whether treatment with antibiotics would lead to the development of a supershedder phenotype, in which an infected host sheds the pathogen at a much higher number than other infected host. This phenomenon has been observed in the natural setting and is considered a major source of host-to-host transmission (48). Murine models have been used to characterize this phenotype, where >10^8^ CFU/g (supershedder [SS] threshold) of the indicated pathogen in the feces is generally considered as the threshold for the supershedder phenotype (SS phenotype) (25, 26). Using the KPPR1S isolate, as it consistently colonized mice at a high density without antibiotic treatment, we assessed fecal shedding of *Kpn* for 10-12 days p.i. after either a single streptomycin treatment or three consecutive days of streptomycin treatment. We found that antibiotic treatment triggered a temporary supershedder phenotype (**Fig. 4A-B**), whereas, no such phenotype was observed with the vehicle only control (PBS) (**Fig. S1A**). A second treatment of antibiotics, after mice had returned to baseline levels of *Kpn* shedding from the first antibiotic treatment, caused the development of another transient supershedder phenotype **(Fig. S1B)**.

**Figure 4.**
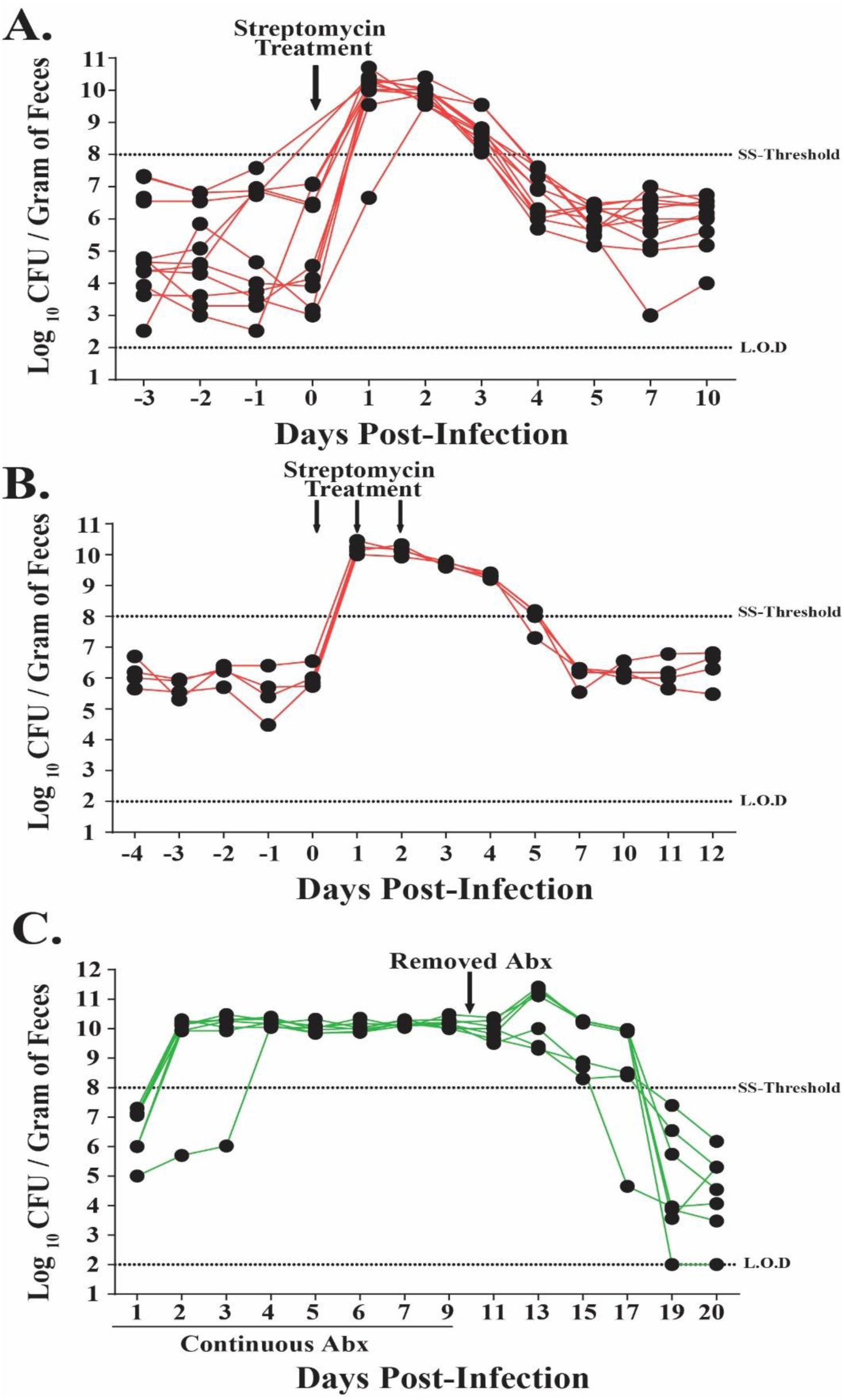
Antibiotic treatment of *K. pneumoniae* infected mice triggers the supershedder phenotype. Fecal shedding from individual mice (n=9) infected with *K. pneumoniae* KPPR1S given a single dose of streptomycin sulfate (5 mg/200 µl) **(A.)** or (n=4) given three treatments of streptomycin on consecutive days **(B.)**, resulted in a rapid development of high shedding (>10^8^ CFU / Gram of feces [Supershedder (SS) threshold] lasting for 3 days post antibiotic treatment. **(C.)** High fecal shedding of *K. pneumoniae* isolate MKP103 from mice (n=6) given ampicillin (1g/Liter) in drinking water. Removal of antibiotic pressure begins an eventual shift to reduced fecal shedding (<10^8^ CFU/Gram of feces).

In a clinical setting, immunocompromised patients tend to be on continuous antibiotic treatment; therefore, we determined the effect of daily antibiotic treatment on *Kpn* shedding. We supplemented the drinking water of mice with ampicillin 24 hours before *Kpn* inoculation and continued for 10 days p.i. As *Kpn* is intrinsically resistant to ampicillin, the mice infected with MKP103 isolate displayed the *Kpn* supershedder phenotype **(Fig. 4C)**. After removal of antibiotic pressure, the mice displayed the high shedding phenotype for multiple days. Taken together, our data suggest that, as a consequence of antibiotic treatment, *Kpn* can develop a supershedder phenotype and the length of this phenotype is dependent upon the duration of the antibiotic treatment.

### Antibiotic treatment leads to the disruption of host-microbiota that correlates with the supershedder phenotype

Next, we determined whether *Kpn* infection or antibiotic treatment induced supershedder phenotype is a result of the displacement of the host microbiota. To provide insight into the *Kpn* carrier state and the supershedder phenotype, we carried out a 16S analysis to determine the host intestinal microbiota changes that occurred during infection and as a consequence of antibiotic treatment (**Fig. 5A**). For a detailed 16S analysis, we isolated DNA from fecal samples collected at six different time points from *Kpn* infected mice (n=4). Fecal pellets were collected pre-inoculation to determine the baseline of the host GI microbiota. Samples were collected on days 7, 9 and 11 post-antibiotic treatment to determine changes in the host microbiota. At days 3 and 5 p.i, we were unable to detect *Kpn* 16S *rRNA* gene sequences, even though it shed at 10^6^ CFU / Gram of fecal sample. This result suggests that *Kpn* comprises only a minor component of the host intestinal microbiota. The main component of a diverse microbial community of the host intestine included *Bacteroidetes* (*Bacteroidales* [*S24*-7]) and *Firmicutes (Clostridales)* **(Fig. 5C; Table. S1)**, which are considered to be a typical profile for stable mammalian intestinal microbiota.

**Figure 5.**
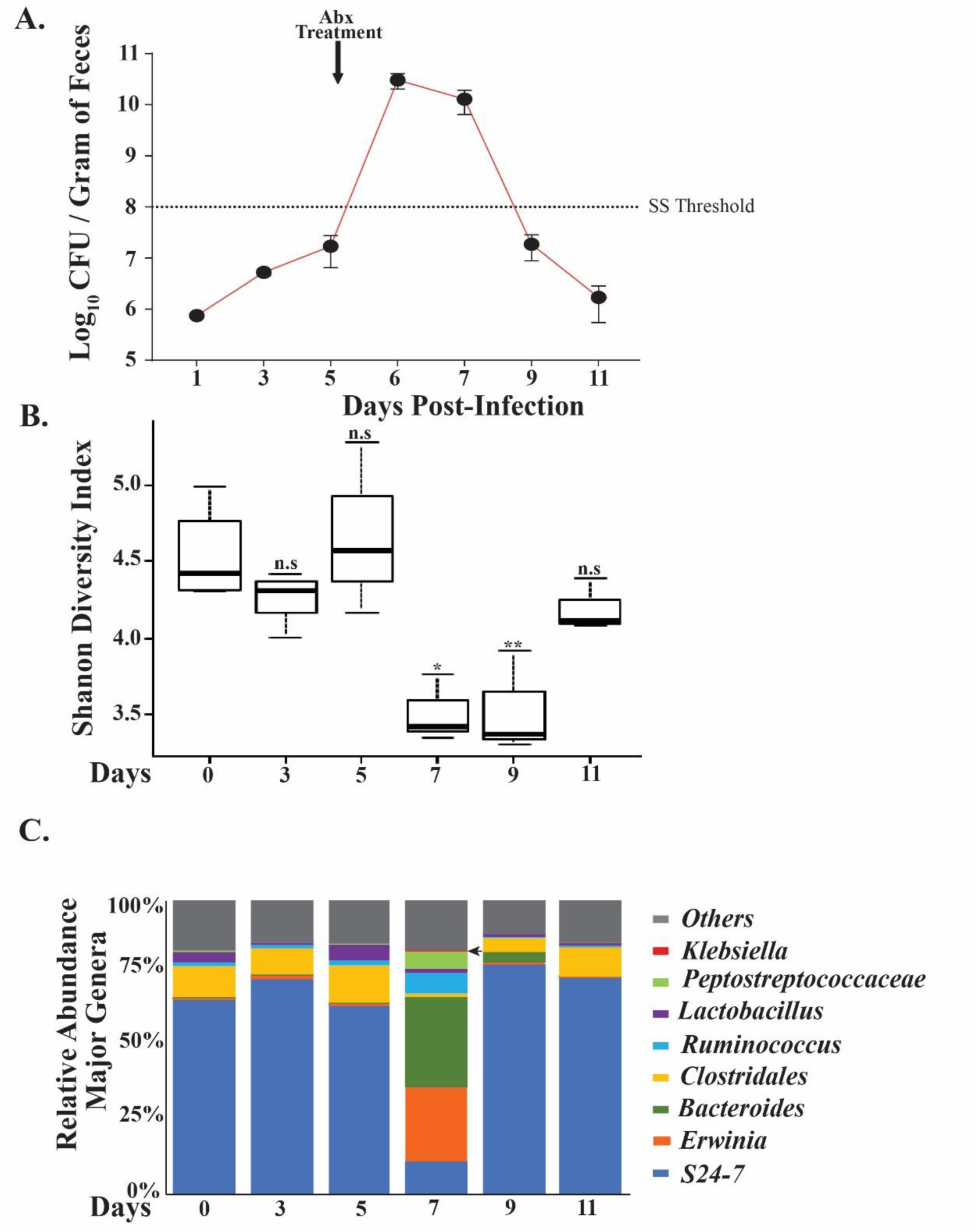
Antibiotic triggered supershedder phenotype correlates with reduced intestinal microbial diversity. **(A.)** Average fecal shedding of *K. pneumoniae* isolate KPPR1S by carrier mice (n=4) pre antibiotic treatment, and post single dose streptomycin treatment (5 mg/200 µl) that induces a transient supershedder phenotype. Error bars represent S.E.M. Dashed line represents the supershedder (SS) threshold. **(B.)** Intestinal microbiota changes in carrier mice from samples obtained on day 0 pre infection, days 3 and 5 post-inoculation, and days 7, 9 and 11 post-antibiotic treatment. DNA was isolated from fecal samples obtained from infected mice and 16s rRNA analysis carried out. Data is shown as Shannon diversity index with mean and *Stdev* values shown. Statistical differences calculated by Kruskal-Wallis test. **(C.)** Shift in microbial diversity determined from fecal samples collected on the days indicated above and shown as bar graph with percent relative abundance of major genera. Arrow shows identification of *Klebsiella pneumoniae* DNA at Day 7 post-infection. *, *p value* < 0.05, **, *p* < *0*.*01*

A single treatment with str led to dramatic changes in the intestinal microbiota. As detailed in **Fig. 5B**, there was a statistically significant decline in the total species richness, especially in *S24-7*, with a concurrent increase in *Erwinia* and *Bacteroides*. As illustrated in **Fig. 5C**, we only observed *Kpn*-specific 16s rRNA gene sequences during the antibiotic-induced supershedder phenotype. A decrease in *Kpn* shedding levels correlated with an increase in *S24-7* and other major components of the host microbiota, and a loss of detection of *Kpn* specific 16s rRNA sequences. Thus, antibiotic treatment leads to a disruption of host-microbiota that correlates with the development of temporary supershedder phenotype. Moreover, disruption of the host microbiota with antibiotics is associated with reduced microbial richness, which recovers three days post-antibiotic exposure.

### *Klebsiella pneumoniae* factors contributing to shedding and colonization

To examine the contribution of known virulence determinants of *Kpn*, we tested shedding and colonization of the previously described capsule (cps)-deficient mutant (Δ*manC*) of the strain KPPR1S. As is evident from **Fig. 6A**, over the course of 15 days of infection, the *ΔmanC* mutant shed and also colonized poorly **(Fig. S2)** in comparison to the parental wild-type (WT) strain.

**Figure 6.**
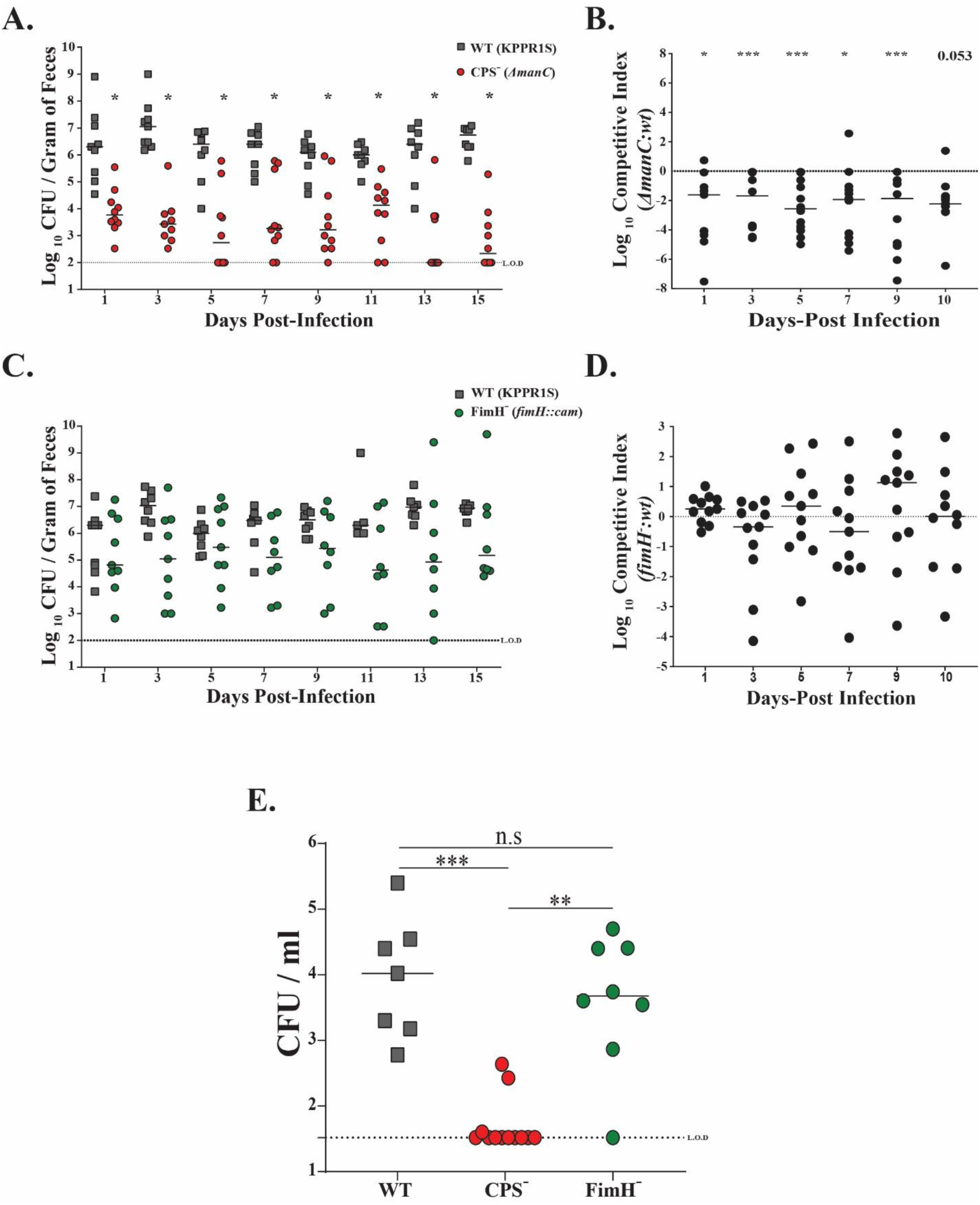
The effect of *K. pneumoniae* virulence factors on fecal shedding and oropharyngeal colonization. **(A.)** Mice were infected orally with KPPR1S (WT) or with a capsule-deficient mutant (*ΔmanC*), and fecal shedding collected on the days indicated. Each symbol represents CFU obtained from a single mouse on a given day, with solid line representing median values. **(B.)** Mice were infected orally with a 1:1 mixture of WT or with capsule-deficient mutant (*ΔmanC*) with fecal shedding collected on the days indicated. The CI was determined as described in the Materials and Methods. Each symbol represents Log^10^ CI value from an individual mouse on a given day. Solid line represents the median. Dashed line indicates a competitive index of 1, or a 1:1 ratio of mutant to WT. **(C.)** Mice were infected orally with WT or with an isogenic mutant (*fimH*::*cam;* FimH-), and fecal shedding collected on the days indicated. **(D.)** Mice were infected orally with a 1:1 mixture of WT or with the mutant (FimH-) with fecal shedding collected on the days indicated. The CI was determined as mentioned in Materials and Methods. **(E.)** The colonization density in the oropharynx for the WT isolate, and isogenic mutants was determined 15 days post-infection with median values shown. For CI statistical differences determined by Wilcoxon signed rank test. Mann-Whitney test used to determine the differences in fecal shedding. Differences in oropharyngeal colonization determined using Kruskal-Wallis test. *, *p* < *0*.*05*, **, *p* < *0*.*01*, ***, *p* < *0*.*001*.

Bacteria can form biofilm like structures in the GI tract (49). We hypothesized that a coinfection with WT *Kpn* and the Δ*manC* strain would form a mixed population (intraspecies) biofilm in the GI tract, helping compensate for the capsule deficiency of the Δ*manC* strain. However, coinfected mice still shed the Δ*manC* strain poorly compared to the parental strain (**Fig. 6B**). These observations suggest that capsular polysaccharide of *Kpn* is essential for robust GI colonization and eventual fecal shedding.

Next, as the type 1 fimbriae of *Kpn* is considered essential for colonization of the host urinary tract, we determined its role in GI colonization (31). The KPPR1S *fim* locus promoter is under phase variable control, which was observed to be in the off position under both *in vitro* (broth culture) and *in vivo* (fecal pellets) (data not shown). To determine the requirement of type 1 fimbriae of KPPR1S in GI colonization, a deletion mutant of *fimH* that encodes the type 1 fimbriae tip adhesin, required for proper interaction with the host epithelial layer (50) was constructed. As is evident from **Fig. 6C-D**, even though mice infected with the *fimH*-mutant had reduced median shedding, it was not significantly lower than the WT strain. Lastly, we determined whether these mutants also contribute towards colonization of the mucosal surface of the oropharynx. **Fig. 6E** shows, capsule was essential for colonization of the oropharyngeal space, whereas type 1 fimbriae was dispensable. Overall our data indicate that *Kpn* capsular polysaccharide plays a critical role in GI colonization. In contrast, *Kpn* type1 fimbriae appears to be nonessential for gut colonization.

### *Klebsiella pneumoniae* transmission occurs through the fecal-oral route

Transmission of enteric pathogens generally occurs through the fecal-oral route, and host-to-host transmission in a hospital setting is a major source of infection (20, 46). Thus, we determined whether *Kpn* host-to-host transmission events could be observed in our animal model. Initially, we housed one uninfected mouse (contact) with four infected mice (index). Fecal pellets were collected to enumerate colonization density and whether transmission from index to contact mice occurred. We observed 100% transmission efficiency with a ratio of 4:1, with transmission occurring within 24 hours of cohousing the animals (**Fig. 7A and C**). Since transmission efficiency is high with a ratio of 4:1, we decided to determine *Kpn* transmission dynamics with one index mouse cohoused with four contact mice. With a ratio of 1:4 reduced transmission efficiency (∼35%) was observed, suggesting that not enough *Kpn* shedding events occurred for all uninfected mice to become colonized (**Fig. 7B-C**).

**Figure 7.**
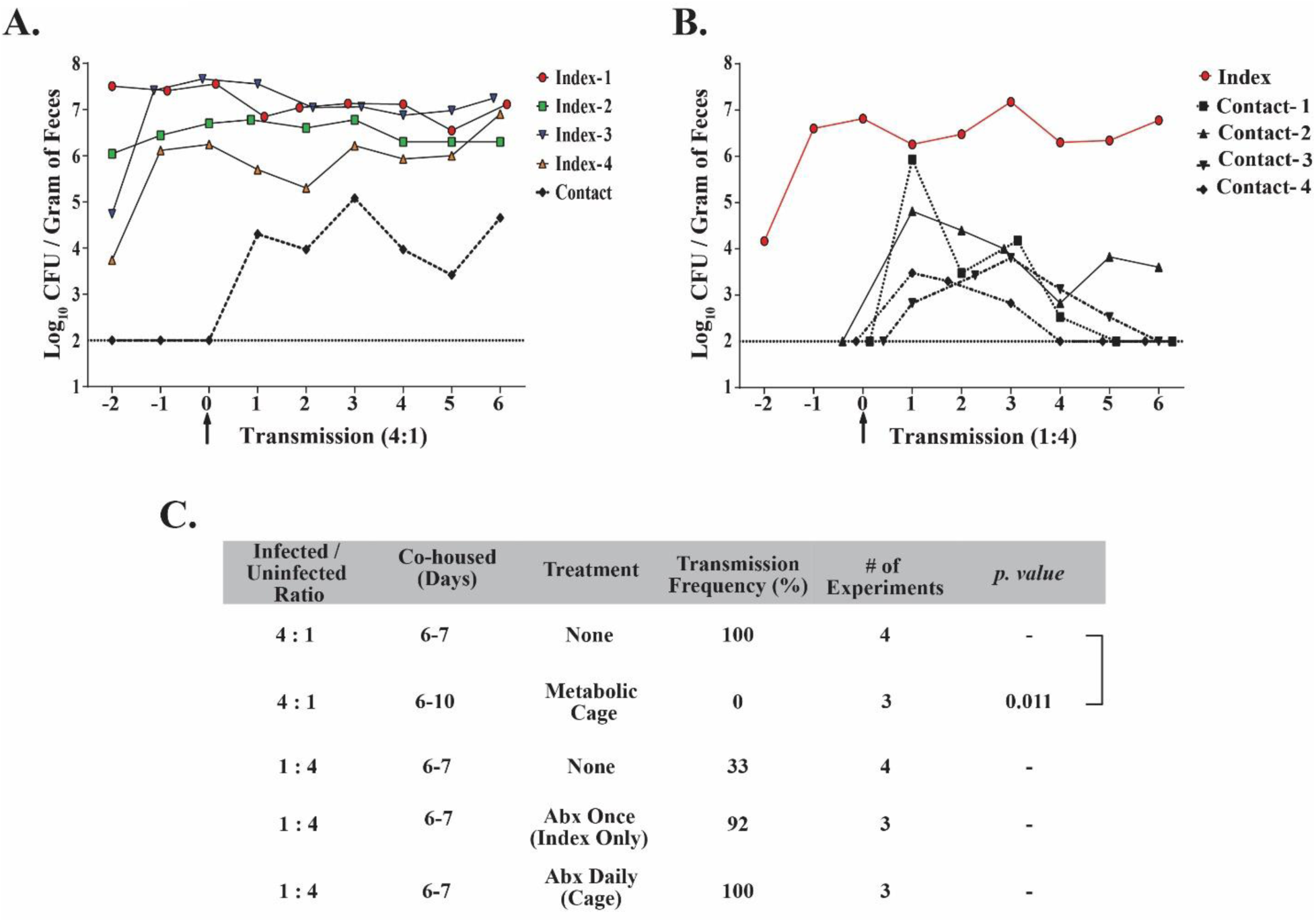
*K. pneumoniae* transmission between hosts with and without treatment of antibiotics. **(A.)** A representative of 4:1 ratio of index to contact transmission. Four naïve mice were infected orally with KPPR1S, and housed with one unifected mouse. Fecal shedding was monitored daily from both the index and the contact mouse. **(B.)** A representative of 1:4 ratio of index to contact transmission. One naïve mouse was infected orally with KPPR1S, and housed with four uninfected mice and fecal shedding monitored daily **(C.)** Observed efficiency of *K. pneumoniae* transmission with different infected to uninfected ratios, effect of antibiotics, and transmission dynamics in a metabolic cage. Statistical differences calculated using two-tailed Fisher’s exact test.

Next, to mimic conditions prevalent in a hospital, where patients tend to be on antibiotics, we investigated the effects of antibiotic treatment on *Kpn* transmission dynamics. A single antibiotic treatment to the index mouse cohoused with four contact mouse led to >90% transmission, suggesting that high *Kpn* shedding in the fecal pellets can overcome colonization resistance of the contact mice (**Fig. 7C**). Lastly, we tested the effect of antibiotics on both index and contact mice by adding antibiotics in their drinking water. An index mouse was infected with MKP103 and was housed separately for several days before being introduced to four contact mice already on antibiotics. We observed 100% transmission efficiency when both index and contact mice were on daily antibiotics. Moreover, as all the mice in the cage were on antibiotics, they all developed the supershedder phenotype (**Fig. S3A**). Our results provide insight into the high transmissibility of *Kpn* in hospitals where there is high antibiotic usage.

We hypothesize that *Kpn* transmission occurs through the fecal-oral route, based upon transmission models of other enteric pathogens and the coprophagic nature of mice. However, as *Kpn* colonizes both the oral cavity and the GI tract, we determined whether host-to-host transmission of *Kpn* is due to the coprophagic action of mice or by contact with infected oral secretions. Mice were housed in a metabolic cage, where they do not have access to their fecal pellets. At a 4:1 ratio of the index to contact mice, no transmission events were detected between the infected and the uninfected mice during the 10-day experiment, suggesting that in our animal model, host-to-host transmission requires the contact with fecal matter (**Fig. 7C**). Lastly, the infected mice in the metabolic cage shed *Kpn* robustly suggesting persistent colonization that did not require re-seeding through the consumption of infected fecal pellets (**Fig. S3B**).

## Discussion

The genetic heterogeneity of *Klebsiella pneumoniae* allows this pathogen to colonize a variety of host mucosal surfaces, which can dramatically impact the clinical outcome. *Klebsiella pneumoniae* disease manifestations in the respiratory and urinary tract have been extensively modeled in animals (4). However, the gastrointestinal mucosal surface, also colonized readily by *Kpn* has not been the focus of many scientific studies (51, 52). In this report, we describe a murine model of oral infection of *K. pneumoniae* to study GI colonization and host-to-host transmission. We demonstrate for the first time that *Kpn* can stably colonize the GI tract of immunocompetent mice without disrupting the host-microbiota – a key strength of our model. Secondly, a host colonized persistently with a pathogen is considered a significant reservoir for new infections, and our animal model of *Kpn* GI colonization replicates this phenotype, indicating it is a useful tool to study within-host events and host-to-host transmission. Third, we observed variability in the ability to colonize the GI tract and to cause invasive disease between different *Kpn* isolates, suggesting *Kpn* genetic plasticity might be involved in the observed variability. Lastly, as many patients in a hospital setting tend to be on antibiotics, we were able to show experimentally for the first time that antibiotic treatment triggers the development of the supershedder phenotype in carrier mice, which promotes host-to-host transmission.

Previous studies used antibiotic treatment to reduce host colonization resistance by disrupting the resident microbiota to establish *Kpn* colonization (31, 41, 53). However, treatment with antibiotics reduces the ability to discern the role of bacterial factors that allow *Kpn* to overcome colonization resistance. Our model does not require the use of antibiotics to establish stable and persistent *Kpn* colonization, and therefore allows for the identification of bacterial and host factors that contribute to *Kpn* colonization and transmission. In our initial studies, we established that the route of infection is critical for stable colonization of *Kpn* in the GI tract. Oral gavage, a standard mode of infection for modeling enteric infections in murine models, only led to transient *Kpn* colonization in the GI tract (data not shown). However, by orally feeding a similar dose allowed *Kpn* to colonize the GI tract and persist without disrupting the host microbiota. A recent study by Atarashi *et al*. showed that *Kpn* colonizing the oral cavity of patients can seed the GI tract (44). In our model of infection we also observed *Kpn* colonizing the murine oral cavity.

A hallmark of many GI pathogens is their ability to cause an acute host inflammatory response. Multiple reports also suggest that *Kpn* might contribute towards gut dysbiosis and play an active role in inducing host response (44, 54). However, epidemiological data also suggest that *Kpn* can silently colonize healthy individuals (18). In our murine model, we were unable to detect any acute signs of inflammation post-*Kpn* infection. Furthermore, unlike the *Salmonella* serovar *Typhimurium* supershedder phenotype, the *Kpn* antibiotic-induced supershedder phenotype was not associated with colitis (**Fig. S4**). Our data suggest that *Kpn* in the GI tract behaves in a manner that does not elicit an acute inflammatory response and carriage is considered an asymptomatic event.

The role of major virulence factors of *Kpn*, including its capsular polysaccharide (CPS), type 1 fimbriae, and others have been extensively examined under both *in vitro* and *in vivo* conditions (4). However, data for the requirement of *Kpn* CPS in GI colonization appears contradictory (41, 42). As those studies were undertaken in mice treated with antibiotics, it is possible that the exact role of the bacterial factors is probably masked. Herein, using our model, we show definitively that CPS of *Kpn* is an essential component required for efficient colonization of both the upper (oropharynx) and lower (intestinal) GI tract. The role of the capsule possibly pertains to protection against host-mediated clearance and interactions with mucus (42, 55, 56).

We also tested the requirement of the *Kpn* type 1 fimbriae in GI colonization. Even though the type 1 fimbriae is critical for colonization of the urinary tract, it appears to be dispensable for GI colonization (31). However, with certain pathogenic *E. coli* isolates, type 1 fimbriae is required for colonization (57, 58). Furthermore, recent work by Jung *et al*. using antibiotic-treated mice observed a defect in GI colonization with a *Kpn fimD* mutant that is missing the usher constituent that facilitates assembly and eventual translocation of the pilus across the outer membrane (53). However, in our model we only observed a slight reduction in median shedding from mice infected with the *fimH* isogenic mutant compared to the WT strain, suggesting that the expression of the type 1 fimbriae of the KPPR1S isolate is dispensable for GI colonization. This result that the type 1 fimbriae of KPPR1S isolate does not contribute towards GI colonization was not unexpected, as we did not observe its expression under the conditions tested (Data not shown). However, type 1 fimbriae might play a role in GI colonization for isolates that do express the structure.

Our model also allows us, for the first time, to understand the transmission dynamics of *Klebsiella pneumoniae*. We observed transmission events between *Kpn*-infected and contact mice, suggesting that *Kpn* shedding in this model is high enough to permit transmission, albeit at a lower frequency. We also observed that the contact mice colonized at a lower density compared to the index mice, possibly because of the reduced *Kpn* dose in the fecal pellets. Thus, our data follows epidemiologic studies that suggest 5-25% carriage rates in the natural environment (19, 59). However, the treatment of either infected or contact mice with antibiotics led to a high host-to-host transmission frequency. A single dose of antibiotic treatment established a suppershedder phenotype in the index host, which was able to transmit to >90% of uninfected mice. It is suggested that 80% of the infections are due to 20% of infected individuals transmitting to uninfected hosts (known as the 80/20 rule) (60). Such individuals are termed as supershedders or superspreaders. However, we were unable to observe a *Kpn* supershedder phenotype without disrupting the host microbiota, suggesting that in our animal model, *Kpn* transmission dynamics do not follow the 80//20 rule. Multiple studies on enteric pathogens show that antibiotic treatment causes a dysbiosis in the GI tract, reduces colonization resistance by the stable resident microbiota, and promotes the expansion of pathogens (25, 26). Our 16S analysis shows that the antibiotic-based supershedder phenotype correlates with a reduction in microbial diversity. In contrast to *Salmonella* serovar *Typhimurium* and *Clostridium difficile* supershedder phenotypes (25, 26), the *Kpn* supershedder phenotype lasts for a shorter duration following a single antibiotic treatment. However, *Kpn* infected mice on continuous antibiotics shed at supershedder levels, a condition we believe to be common in a hospital setting. Our data suggest that the development of the supershedder phenotype is the main contributor to host-to-host transmission events in a hospital. The transmission frequency of *K. pneumoniae* has not been established in a hospital setting and may be higher or lower than the rates determined in our murine model. We believe that a setting with high antibiotic use increases the likelihood of *Kpn* outbreaks. Therefore, patients on antibiotics should be carefully monitored to determine if they are colonized with *K. pneumoniae*.

In conclusion, we have described a model that will be useful in understanding complex interactions between *K. pneumoniae* and the host immune system and the intestinal microbiota. The availability of an arrayed marked mutant library of *Kpn* (32) and several annotated *Kpn* genomes should allow for studies identifying bacterial factors that contribute towards *Kpn* colonization and transmission. Since a majority of *Kpn* nosocomial infections arise from GI colonization and fecal-oral route of transmission (20, 29), an understanding of the biology of *Kpn* gastrointestinal colonization and fecal-oral transmission would be valuable as it could serve as an ideal point of intervention.

## Acknowledgements

We would like to thank Drs. Michael Bachman (University of Michigan), Alan Hauser (Northwestern University), Virginia Miller (UNC Chapel Hill), Thomas Russo (University of Buffalo-SUNY) and Jeffery N. Weiser (NYU School of Medicine) for the strains used in this study. We would also like to thank Drs. Phillip Hernandez (Boston University), Virginia Miller, Kimberly Walker (UNC Chapel Hill), Jeffrey N. Weiser and Tonia Zangari (NYU School of Medicine) for fruitful discussions in regards to the establishment of the model and the manuscript.

This study was funded by startup funds provided by Wake Forest Baptist Medical Center to M.A.Z.

## Supplemental Figures

**Figure S1.**
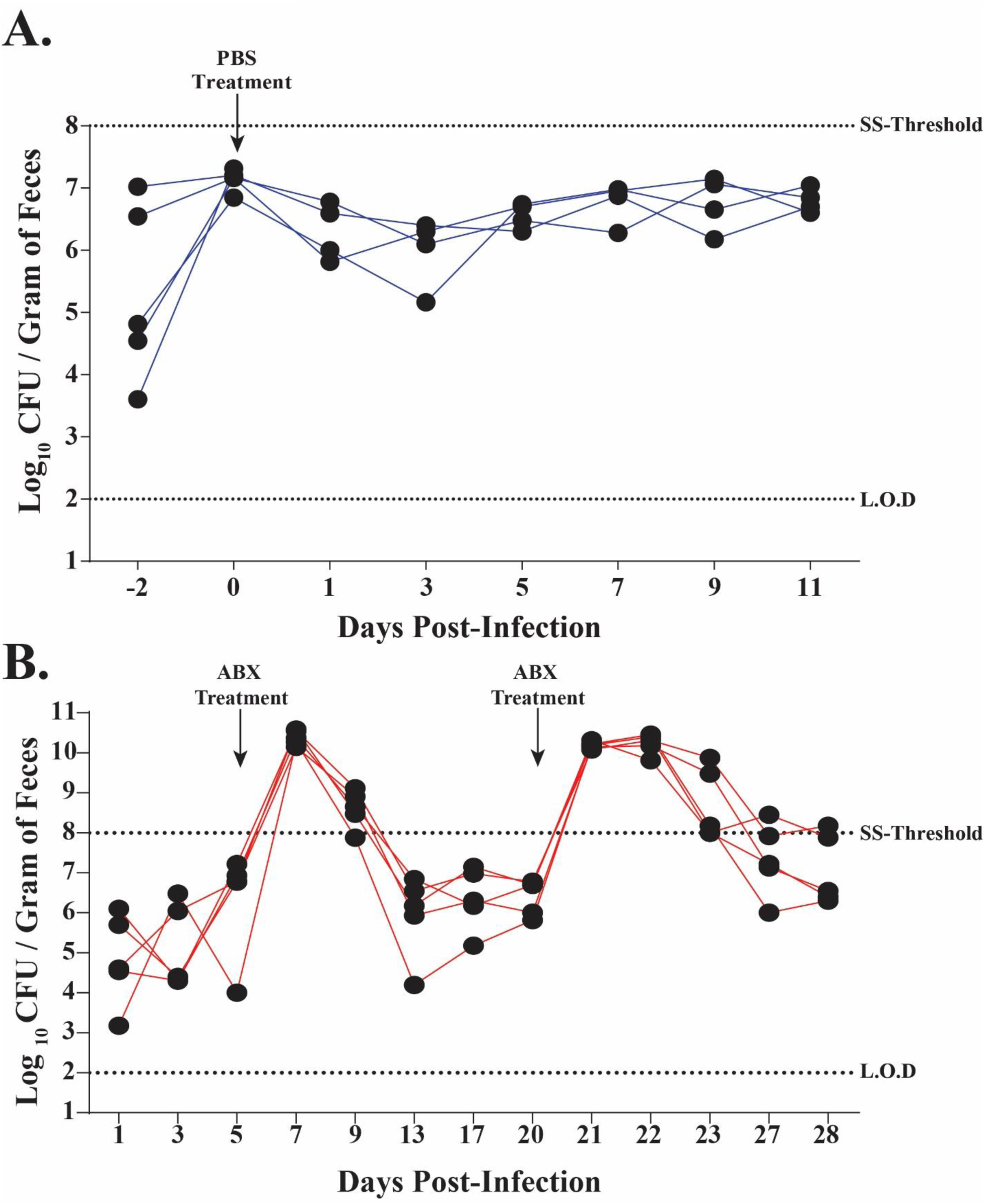
**(A.)** Treatment of *K. pneumoniae* infected mice with PBS alone (200 µl) does not elicit a supershedder phenotype. **(B.)** Streptomycin sulfate (5 mg/200 µl) treatment triggers the supershedder phenotype (> 10^8^ CFU/Gram of feces) in mice infected with *Klebsiella pneumoniae*. A second treatment of Streptomycin sulfate (5 mg/200 µl) once mice have returned to reduced *Kpn* shedding elicits a second supershedder phenotype that is also transient.

**Figure S2.**
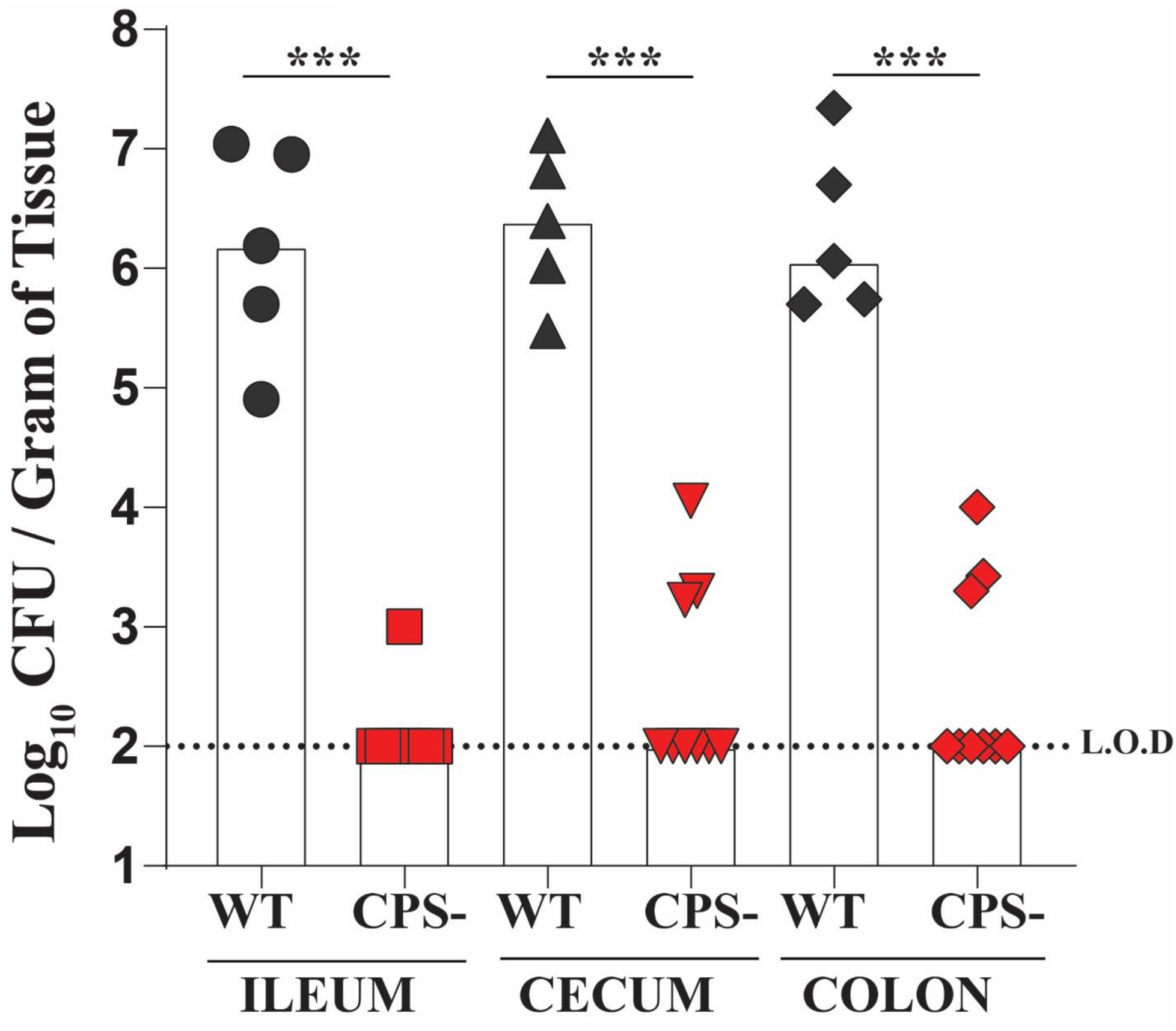
The colonization density in the GI tract (Ileum, cecum and colon) for the WT isolate, and isogenic capsule deficient mutant (*ΔmanC*) was determined 15 days post-infection with median values shown. Limit of detection was 10^2^ CFU/Gram of tissue. Differences in GI colonization determined using Kruskal-Wallis test. ***, *p* < *0*.*001*.

**Figure S3.**
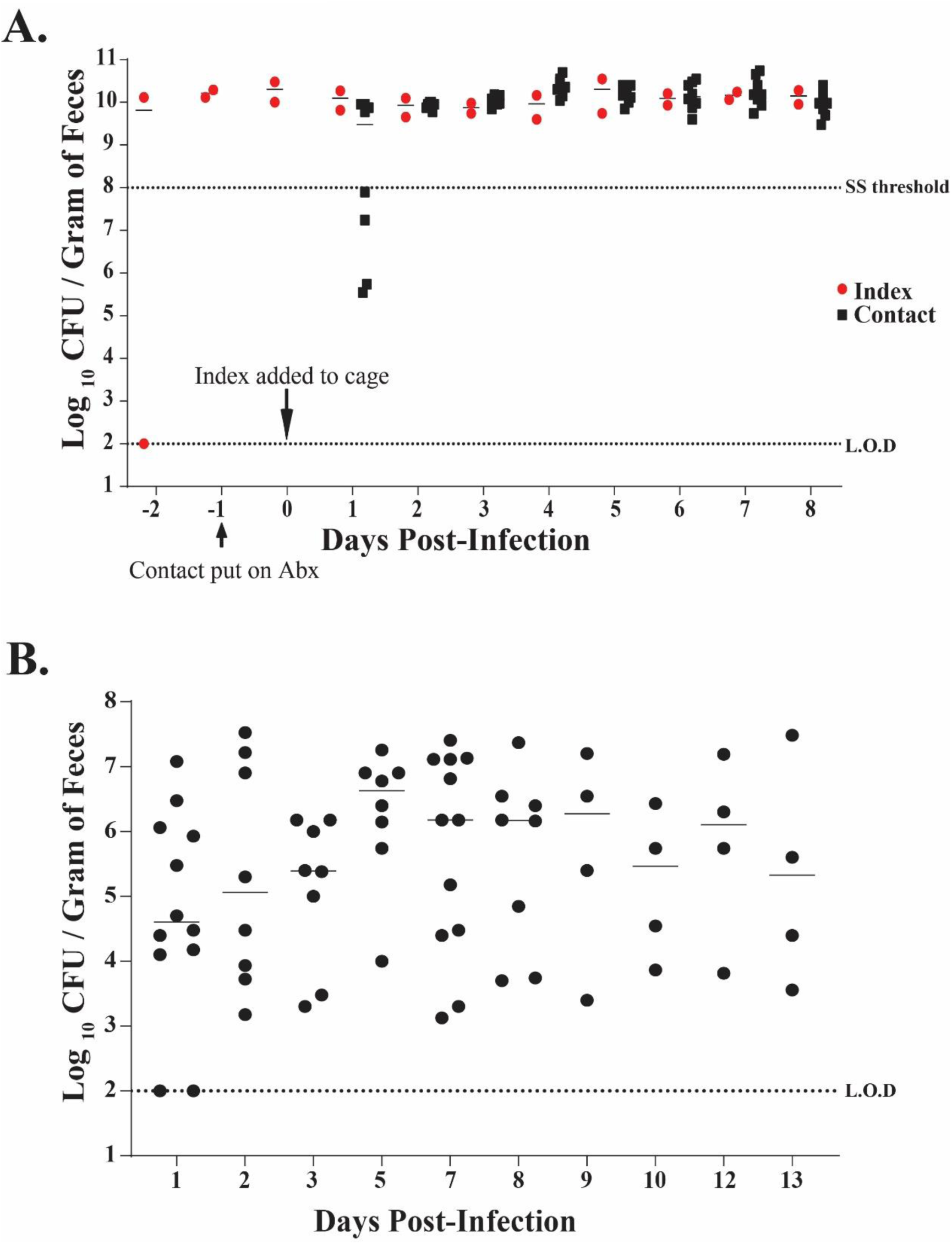
**(A.)** Antibiotic treatment leads to development of supershedder phenotype in the index (Infected) mice and high transmission rates. Index mice were given ampicillin (1g/Liter) in drinking water a day before oral inoculation with *Kpn* isolate MKP103. One day before the index mouse was moved in to the contact mice (n=4) cage they were also given ampicillin (1g/Liter) in drinking water. The contact mice quickly became colonized with MKP103 isolate, and both the index and contact mice displayed the supershedder phenotype (> 10^8^ CFU/Gram of feces). Shown here are representation of two independent transmission studies. **(B.)** Fecal shedding data collected from mice given 10^6^ CFU of KPPR1S housed in a metabolic cage to reduce coprophagia, and followed for up to either 13 days post-infection. Shown are median values, where each point represents a single mouse on a given day. Limit of detection (L.O.D) was 10^2^ CFU/Gram of Feces.

**Figure S4.**
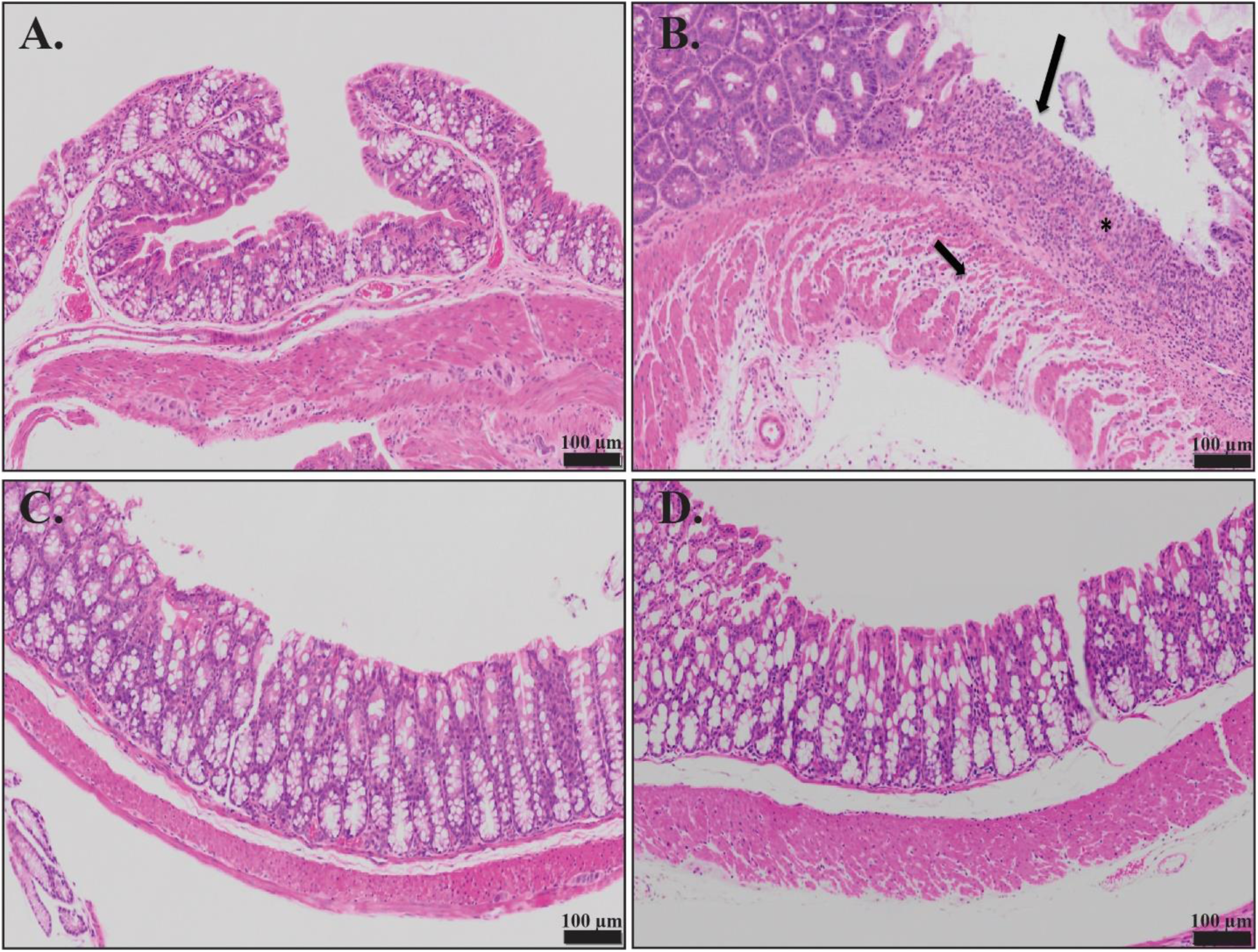
Colonization by *Klebsiella pneumoniae* shows normal colonic mucosa. Displayed here are swiss rolls of colonic mucosa stained with hematoxylin and eosin staining. **(A.)** Mice given PBS alone and tissue prepared 7 days post-treatment. **(B.)** Colon tissue from mice prepared 7 days post 3% DSS daily treatment. Detailed here is the ulceration (long arrows) of the mucosa, with inflammation expanding the submucosa (*), and loss and fragmentation of individual muscle cells (short arrows). **(C.)** Mice infected with KPPR1S and tissue prepared 7 days post-infection. **(D.)** Mice infected with KPPR1S, given a single-dose of streptomycin (5 mg/ 200 µl) to trigger supershedder phenotype and tissue prepared two days post antibiotic treatment. Colonic epithelium of mice appears normal that are infected with KPPR1S and its supershedder phenotype, and is similar in morphology to the PBS control.

**Table S1.**
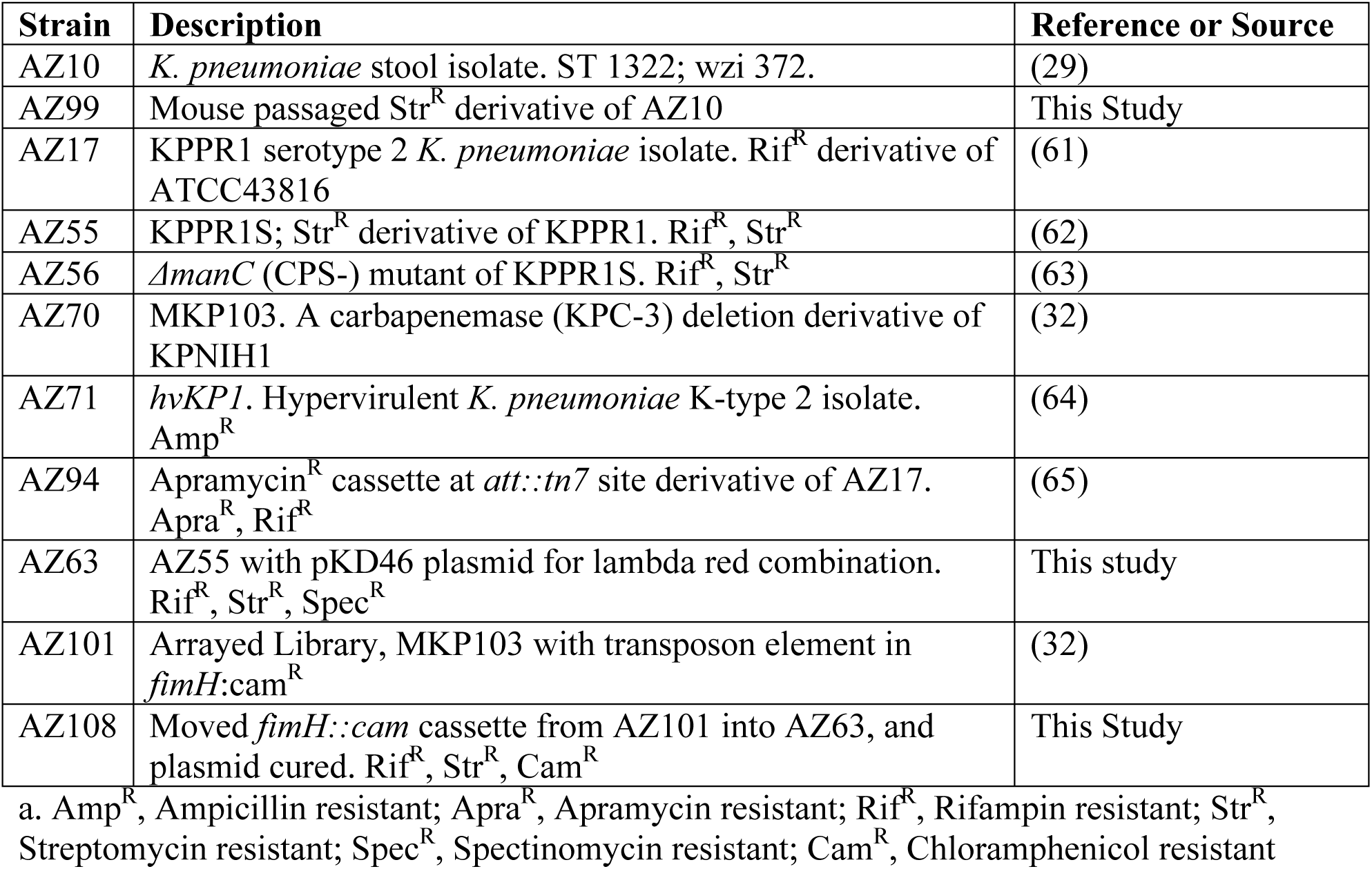
List of bacterial species identified in the mice fecal pellets using 16s rRNA analysis Strains used in this study

